# Identification of levoglucosan degradation pathways in bacteria and sequence similarity network analysis

**DOI:** 10.1101/2023.01.28.526070

**Authors:** Arashdeep Kaur, Nichollas E. Scott, Marion Herisse, Ethan D. Goddard-Borger, Sacha Pidot, Spencer J. Williams

## Abstract

Levoglucosan is produced in the pyrolysis of cellulose and starch, including from bushfires or the burning of biofuels, and is deposited from the atmosphere across the surface of the earth. We describe two levoglucosan degrading *Paenarthrobacter* spp. (*Paenarthrobacter nitrojuajacolis* LG01 and *Paenarthrobacter histidinolovorans* LG02) that were isolated from soil by metabolic enrichment using levoglucosan as the sole carbon source. Genome sequencing and proteomics analysis revealed the expression of a series of genes encoding known levoglucosan degrading enzymes, levoglucosan dehydrogenase (LGDH, LgdA), 3-keto-levoglucosan β-eliminase (LgdB1) and glucose 3-dehydrogenase (LgdC), along with an ABC transporter cassette and an associated solute binding protein. However, no homologues of 3-ketoglucose dehydratase (LgdB2) were evident, while the expressed genes contained a range of putative sugar phosphate isomerases/xylose isomerases with weak similarity to LgdB2. Sequence similarity network analysis of genome neighbours of LgdA revealed that homologues of LgdB1 and LgdC are generally conserved in a range of bacteria in the phyla Firmicutes, Actinobacteria and Proteobacteria. One group of sugar phosphate isomerase/xylose isomerase homologues (named LgdB3) was identified with limited distribution that is mutually exclusive with LgdB2, and we propose that they may fulfill a similar function. LgdB1, LgdB2 and LgdB3 adopt similar predicted 3D folds, suggesting overlapping function in processing intermediates in LG metabolism. Our findings highlight diversity within the LGDH pathway, through which bacteria utilize levoglucosan as a nutrient source.

## Introduction

Levoglucosan (LG; 1,6-anhydro-β-D-glucose) is a bicyclic anhydride of D-glucose (McGill and Williams 2008). It is formed by a range of chemical processes including pyrolysis of glucose (Sugisawa and Edo 1966), starch and cellulose (Lakshmanan et al. 1969; Itabaiana Junior et al. 2020). LG is formed from the burning of cellulose above 300 °C (Simoneit et al. 1999) and is produced by the burning of biomass (Suciu et al. 2019). Emissions of LG into the atmosphere arise from burning of plant matter in open fires (1.7 million tonnes per annum) and biofuels (2.1 million tonnes per annum) and contributes to aerosols in the troposphere that affect air quality (Li et al. 2021). Consequently, LG can be used as a molecular marker for biomass burning and as a measure of air quality (Li et al. 2021), and has been used to trace ancient fire histories (paleo-fire records) in glacier snow and ice (You and Xu 2018). While LG is short lived in the atmosphere and degrades with a lifetime of 1.8 days, an estimated 0.7 million tonnes of LG are deposited annually through dry deposition and in rain: ‘sugar from the heavens’ (Li et al. 2021). Consequently, soils across the earth likely experience continuous, low levels of LG deposition, irrespective of proximity to biomass combustion processes or precipitation.

Microorganisms that can grow on LG as sole carbon source are readily recovered from soils (Kitamura et al. 1991; Yasui et al. 1991; Nakahara et al. 1994; Iwazaki et al. 2018) including those associated with wood burning (Arya et al. 2022), as well as plant leaves (Iwazaki et al. 2018), and pond (Iwazaki et al. 2018) and waste water (Arya et al. 2022), suggesting that LG degrading organisms are widespread in Nature. While humans and mice do not metabolize LG (Migliaccio et al. 2009; Bergauff et al. 2010), pathways for its degradation have been elucidated in fungi and bacteria. Fungi appear to exclusively use a pathway involving LG kinase, an ATP-dependent enzyme that converts LG in one step to glucose-6-phosphate (Kitamura and Yasui 1991; Bacik et al. 2015). On the other hand, bacteria use a multistep pathway (Yasui et al. 1991) that was first elucidated in *Bacillus smithii* S-720M (Kuritani et al. 2020) (**Figure 1**). In this organism, LG metabolism is initiated by the action of NADH-dependent LG dehydrogenase (LGDH, also known as LgdA), which forms 3-keto-LG. LGDH from *Pseudarthrobacter phenanthrenivorans* Sphe3 has been studied extensively, including determination of its 3D structure, identification of its active site, and the binding mode of LG (Sugiura et al. 2018). The 1,6-anhydro ring of 3-keto-LG is cleaved to give 2-hydroxy-3-keto-d-glucal, catalyzed by 3-keto-LG β-eliminase (LgdB1), and this is hydrated to give 3-keto-d-glucose, catalysed by 3-ketoglucose dehydratase (LgdB2). Finally, 3-keto-d-glucose is reduced to D-glucose, catalyzed by NADH-dependent glucose 3-dehydrogenase (LgdC). LgdB2 is a bifunctional enzyme that can also catalyze the conversion of 3-keto-LG to 2-hydroxy-3-keto-d-glucal directly. Overall, this complex pathway involves 3-4 enzymes to achieve the formal hydrolysis of LG. This pathway bears some similarity to glycoside hydrolases of carbohydrate active enzyme family GH4 (Yip et al. 2004) and GH109 (Liu et al. 2007), which in a single step catalyze glycoside hydrolysis in an NADH-dependent manner through a redox mechanism involving a 3-keto sugar intermediate (Yip and Withers 2006), and the 2,7-anhydro-N-acetylneuraminic acid hydrolase from *Ruminococcus gnavus*, which cleaves this anhydro sugar via a 4-keto sialic acid intermediate (Bell et al. 2020). Possibly, the requirement of multiple enzymes for the net hydrolysis of LG relates to the profound conformational change that occurs when the 1,6-anhydro ring of LG is cleaved, meaning that different enzymes are required to recognize the substrate and intermediates in the pathway.

**Fig. 1.**
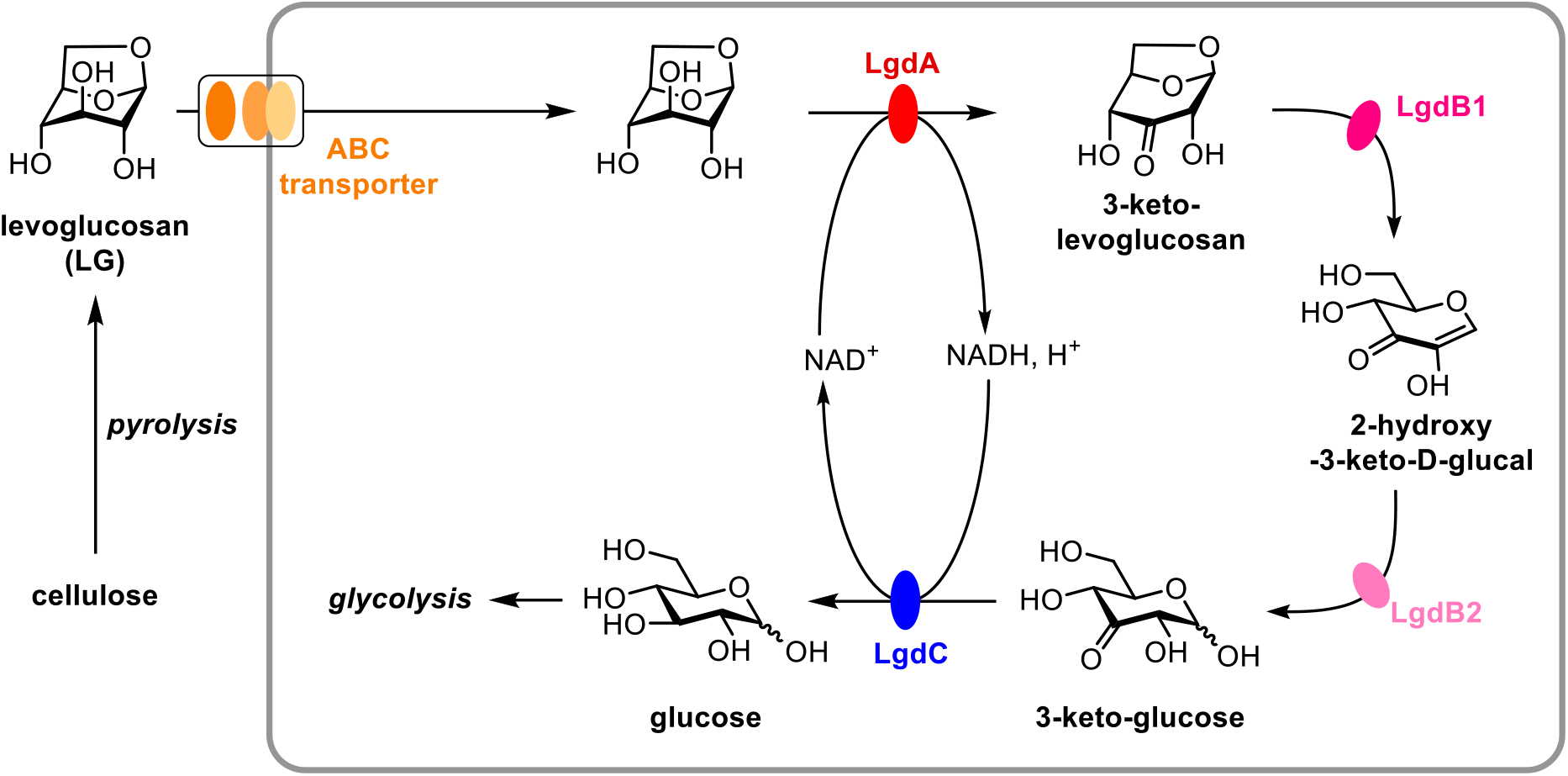
Production of levoglucosan (LG) by pyrolysis of cellulose, and importation and metabolism of LG in *Bacillus smithii* S-2701M. NADH-dependent LG dehydrogenase (LgdA, also known as LGDH) oxidizes LG to 3-keto-LG. 3-Keto-LG β-eliminase (LgdB1) cleaves the 1,6-anhydro ring to give 2-hydroxy-3-keto-d-glucal. 3-Keto-glucose dehydratase (LgdB2) catalyses the hydration of 2-hydroxy-3-keto-d-glucal. Glucose 3-dehydrogenase (LgdC) catalyzes the reduction of 3-keto-d-glucose to D-glucose. LgdB2 is reported to be a bifunctional enzyme that can also catalyze the conversion of 3-keto-LG to 2-hydroxy-3-keto-d-glucal.

All LG-degrading bacteria studied at a genetic level contain close homologs of *lgdA*, and previous work has sought to identify genes involved in LG transport and metabolism by analysis of the surrounding genomic regions. In the case of *Bacillus smithii* S-270M, a genomic region containing *lgdA-lgdB1-lgdB2-lgdC* also contains genes encoding ABC transporter subunits and an ABC solute binding protein (Kuritani et al. 2020). However, other LG degrading organisms such as *P. phenanthrenivorans* Sphe3, *Paenibacillus athensensis* MEC069, *Shinella sumterensis* MEC087, *Microbacterium* MEC084, and *Klebsiella pneumoniae* MEC097 contain *lgdA-lgdB1* and genes encoding ABC transporter subunits and substrate binding proteins, but homologs of *lgdB2* and *lgdC* were not identified (Arya et al. 2022), raising questions as to whether other enzymes may be used for the final steps of LG metabolism.

Here, we isolate two LG degrading bacteria from soil and report their draft genome sequences. Both organisms contain close homologs of *lgdA* and *lgdC* that encode the enzymes that catalyze the first and last steps of the bacterial LG degradation pathway, but lack the full complement of homologues for the intervening genes. We perform comparative proteomics to identify the genes that are upregulated upon growth on LG, and use sequence similarity network analysis to study the genome neighbourhoods surrounding *lgdA*. We identify a range of putative sugar phosphate isomerase/xylose isomerases with weak similarity to LgdB2 that have a distribution that is mutually exclusive with LgdB2, and which we suggest may fulfil a similar function. Our findings highlight the diversity of bacterial LGDH pathways through which bacteria utilize levoglucosan as a nutrient source.

## Materials and methods

### Bacterial growth media

LG growth media was prepared using M9 salt medium, trace metal solution and vitamin mixture and 5 mM levoglucosan as sole carbon source. M9 minimal medium consisted of 12.8 g/L Na_2_HPO_4_, 3 g/L KH_2_PO_4_, 1 g/L NH_4_Cl, 0.5 g/L NaCl, 0.6 g/L MgSO_4_, 0.22 g/L CaCl_2_.2H_2_O salts and trace metal solution consisted of 14.4 mg/L FeCl_3_, 4.0 mg/L CoCl_2_.6H_2_O, 2.9 mg/L CuCl_2_.2H_2_O, 3.0 mg/L MnCl_2_.4H_2_O, 15.0 mg/L ZnCl_2_, 0.2 mg/L NiCl_2_, 3 mg/L Na_2_MoO_4_.6H_2_O and 6 mg/L H_3_BO_3_. Vitamin mixture consisted of 2 μg/L biotin, 5 μg/L calcium pantothenate, 10 μg/L thiamine hydrochloride, 10 μg/L p-aminobenzoic acid, 20 μg/L nicotinic acid, 50 μg/L pyridoxamine dihydrochloride, and 4 μg/L B12 (cyanocobalamin).

#### Isolation of levoglucosan degrading species

*Paenarthrobacter nitrojuajacolis* LG01 and *Paenarthrobacter histidinolovorans* LG02 were isolated from enriched culture, obtained from granular soil sampled from the A horizon, Royal Park (Melbourne, Australia). There was no apparent fire history of the sampled soil.

Approximately 1 g of soil samples were suspended in 5 mL of sterilized LG growth media containing 5 mM of LG as sole carbon source. The culture was incubated at 28 °C for 5 days with continuous shaking at 250 rpm. A subsample of 50 μL was supplemented into freshly prepared levoglucosan media and grown for a further 3 days. This step was repeated 5 times and after the 5^th^ transfer, cells were plated on LB agar plates consisting of 10 g/L tryptone, 5 g/L NaCl, 5 g/L yeast, 15 g/L agar and incubated overnight at 28 °C in the dark. Single colonies were picked and used to inoculate fresh LG growth media and incubated at 28 °C with continuous shaking at 250 rpm. Then, cells were plated and re-inoculated into fresh liquid LG growth media 2 more times. Stock cultures were prepared by addition of 10% glycerol and stored at −80 °C. Genomic DNA of LG degrading species was extracted using the GenElute DNA extraction kit (Sigma) with inclusion of lysozyme and RNAase.

### Phenotypic assays

Strains LG01 and LG02 were grown to visible turbidity in LG growth media containing 20 mM of levoglucosan at 28 °C for 3 days with continuous shaking at 250 rpm. Cultures were centrifuged at 10000 rpm for 10 min (using Sigma laborzentrifugan model 1-15, rotor 12124) and the supernatant liquid was diluted with 50% D_2_O. ^13^C-NMR spectra of diluted supernatant liquid was recorded using a 500 MHz instrument.

### Genome sequence, assembly, and annotation

Genomic DNA was sequenced using an Illumina MiSeq at the Peter Doherty Institute for Infection and Immunity, Parkville, Victoria, Australia. DNA was prepared for sequencing on the Illumina MiSeq platform using the Nextera XT DNA preparation kit (Illumina) with ×300 bp paired end chemistry and with a targeted sequencing depth of >50×. Draft genomes were assembled using Shovill v1.1.0 (https://github.com/tseemann/shovill) and annotated using Prokka v1.14.5 (Seemann 2014). GC percentage and ANI calculations were performed using ANI calculator (https://www.ezbiocloud.net/tools/ani) (Yoon et al. 2017). Taxonomic annotations used 16S rRNA sequences extracted from the assembled genomes as a query in a BLASTn search against the NCBI 16S rRNA database. Assembled genomes have been deposited at the NCBI (Bioproject accession: LG01, PRJNA925556; LG02, PRJNA925557). General features for isolated bacteria are reported in Table 1.

**Table 1.** Key features of the LG01 (BioProject accession: PRJNA925556) and LG02 (BioProject accession: PRJNA925557) draft genome assemblies.

|  | **LG01** | **LG02** |
| --- | --- | --- |
| Genome size | 4777618 | 4332355 |
| Number of contigs | 86 | 862 |
| GC % | 62.86 | 62.92 |
| Number of ORFs | 4455 | 4403 |
| Number of putative genes | 4519 | 4462 |
| Number of putative tRNA | 56 | 55 |
| Number of putative rRNA | 7 | 3 |
| Number of putative tmRNA | 1 | 1 |
| Number of genes assigned a function (%) | 2419 (53%) | 2094 (47%) |

### Sample preparation for proteomic analysis

Cells were grown to mid log phase and washed 3 times with phosphate buffered saline (PBS), collected by centrifugation at 10000 rpm, and then snap frozen in liquid nitrogen and stored at −80 °C. Pelleted snap frozen bacterial cells were solubilized in 4% SDS, 100 mM HEPES by boiling for 10 min at 95 °C. Protein concentrations were assessed using the bicinchoninic acid protein assay (Thermo Fisher Scientific) and 100 μg of each independent replicate prepared for digestion using Micro S-traps (Protifi, USA) according to the manufacturer’s instructions. Samples were reduced with 10 mM DTT for 10 mins at 95 °C and then alkylated with 40 mM iodoacetamide in the dark for 1 h. Samples were acidified to 1.2% phosphoric acid and diluted with seven volumes of S-trap wash buffer (90% methanol, 100 mM tetraethylammonium bromide, pH 7.1), then were loaded onto S-traps and washed 3 times with 400 μl of S-trap wash buffer. Samples were then digested with 3 μg of trypsin (a 1:33 protease/protein ratio) in 100 mM tetraethylammonium bromide for 4 h at 37 °C, then were collected by centrifugation with washes of 100 mM tetraethylammonium bromide, followed by 0.2% formic acid followed by 0.2% formic acid / 50% acetonitrile. Samples were dried and further cleaned up using C_18_ Stagetips to ensure the removal of particulate matter (Rappsilber et al. 2003; Rappsilber et al. 2007).

### Reverse phase liquid chromatography–mass spectrometry for proteomics

Prepared purified peptides from each sample were re-suspended in Buffer A* (2% acetonitrile, 0.01% trifluoroacetic acid) and separated using a two-column chromatography setup composed of a PepMap100 C_18_ 20-mm by 75-μm trap and a PepMap C_18_ 500-mm by 75-μm analytical column (Thermo Fisher Scientific). Samples were concentrated onto the trap column at 5 μl/min for 5 min with Buffer A (0.1% formic acid, 2% DMSO) and then infused into an Orbitrap Eclipse Mass Spectrometer (Thermo Fisher Scientific) or a Orbitrap Fusion Lumos equipped with a FAIMS Pro interface both at 300 nl/minute via the analytical columns using a Dionex Ultimate 3000 UPLC (Thermo Fisher Scientific). For samples infused into the Orbitrap Fusion Lumos, 125-minute analytical runs were undertaken by altering the buffer composition from 2% Buffer B (0.1% formic acid, 77.9% acetonitrile, 2% DMSO) to 22% B over 95 min, then from 22% B to 40% B over 10 min, then from 40% B to 80% B over 5 min. The composition was held at 80% B for 5 min, and then dropped to 2% B over 2 min, then held at 2% B for another 8 min. For samples infused into the Orbitrap Eclipse 95-minute analytical runs were undertaken by altering the buffer composition from 2% Buffer B (0.1% formic acid, 77.9% acetonitrile, 2% DMSO) to 22% B over 65 min, then from 22% B to 40% B over 10 min, then from 40% B to 80% B over 5 min. The composition was held at 80% B for 5 min, and then dropped to 2% B over 2 min, then held at 2% B for another 8 min. The Orbitrap Eclipse mass spectrometer was operated in a data-dependent mode automatically switching between the acquisition of a single Orbitrap MS scan (375-1500 *m/z* and a resolution of 120k) and 3 seconds of Orbitrap MS/MS HCD scans of precursors (NCE of 30%, a maximal injection time of 22 ms, and a resolution of 15k). The Fusion Lumos Mass Spectrometer was operated in a stepped FAIMS data-dependent mode at two different FAIMS CVs −40 and −60. For each FAIMS CV a single Orbitrap MS scan (300-1600 m/z and a resolution of 60k) was acquired every 1.5 seconds followed by Orbitrap MS/MS HCD scans of precursors (Stepped NCE 28,32,37%, maximal injection time of 100 ms and a resolution of 30k).

### Proteomic data analysis

Identification and LFQ analysis were accomplished using MaxQuant (v2.0.2.0) (Cox and Mann 2008) using the in house generated proteomes of LG01 and LG02 allowing for oxidation on Methionine. Prior to MaxQuant analysis datasets acquired on the Fusion Lumos were separated into individual FAIMS fractions using the FAIMS MzXML Generator (Hebert et al. 2018). The LFQ and “Match Between Run” options were enabled to allow comparison between samples. The resulting data files were processed using Perseus (v1.4.0.6) (Tyanova et al. 2016) to compare the growth conditions using Student’s t-tests as well as Pearson correlation analyses.

### Bioinformatic analysis

The protein sequences of LG degrading enzymes from *Pseudarthrobacter phenanthrenivorans* Sphe3 and *B. smithii* S-270M were used as queries to search the genomes of LG01 and LG02 using BLASTp.

The LGDH sequence from isolated strain LG01 was submitted as query to phmmer web server (https://www.ebi.ac.uk/Tools/hmmer/search/phmmer) (Potter et al. 2018). Standard parameters were used except the cut-off for *E*-value was set to 10^−130^. The resulting sequences were used to build a hidden Markov model (HMM), which was used as query for *hmmsearch* in hmmer web server (https://www.ebi.ac.uk/Tools/hmmer/search/hmmsearch). The results were filtered and protein sequences with *E*-value ≤ −200 (**see Supplementary file 2**) were used to generate a sequence similarity network (SSN) using the Enzyme Function Initiative-Enzyme Similarity Tool (https://efi.igb.illinois.edu/efi-est/) with minimum alignment score 120 (corresponding to ≥60% identity) and 170 (corresponding to ≥70% identity; **see Supplementary file 3**) (Zallot et al. 2019). The SSN generated using the alignment score 120 was used to generate genome neighbourhood diagrams (GND) with open reading frame (ORF) ± 10 neighbours using the Enzyme Function Initiative-Genome Neighbourhood Tool (https://efi.igb.illinois.edu/efi-gnt/). A script was used to extract the accession codes of the retrieved neighbours (**Figure S1**). The LGDH neighbours were used to generate an SSN of neighbours (SSNN; **see Supplementary file 4**). The resulting SSNN was plotted using Cytoscape v3.9, with minimum alignment score of 50.

The individual nodes of the LGDH SSN were manually coloured according to the occurrence of *lgdB1, lgdB2, lgdB3* and *lgdC* gene homologues in their neighbourhood (ORF ± 10 neighbours). Initially, the EFI tool was used to generate a cluster number for each unique isofunctional cluster within the LGDH SSNN. The cluster number of the neighbourhood genes from the SSNN for *lgdB1, lgdB2, lgdB3* and *lgdC* were mapped onto the LGDH SSN (alignment score of 170).

### Structural analysis of LG degrading proteins

3D structural models were predicted for LgdA, LgdB1, LgdB2, LgdB4, LgdB4 and LgdC proteins using AlphaFold2 (Jumper et al. 2021; Varadi et al. 2022).

## Results

### Isolation and characterization of bacteria that can grow on LG

Bacteria able to grow on LG were selected by using soil samples (collected from the South-East corner of Royal Park, Melbourne) to inoculate a vitamin-enriched minimal media containing LG as sole carbon source. Sequential subculturing into fresh LG-minimal media, followed by repeated plating onto LB agar and then regrowth in LG-minimal media, led to isolation of strains LG01 and LG02 (**Table S1**). Strains LG01 and LG02 grew robustly on LG with peak growth rates of 0.073 and 0.288 A_600_/min and achieved stationary phase after approximately 82 and 18 h, respectively (**Figure S2**). The absorbance at stationary phase for cultures grown on LG were approximately the same for cultures grown on equimolar glucose. ^13^C-NMR analysis of spent culture medium of LG01 and LG02 grown on LG (20 mM) media did not reveal any carbon containing metabolic products.

To investigate the genetic basis for LG consumption by LG01 and LG02, genomic DNA extracted from these bacteria were sequenced using the Illumina NextSeq platform. **Table 1** shows the key features of the two draft genomes. Based on 16S rRNA gene sequence analysis the organisms were assigned as *Paenarthrobacter nitrojuajacolis* LG01 and *Paenarthrobacter histidinolovorans* LG02.

### Identification of genetic loci expressed upon growth on LG

To identify possible proteins involved in LG metabolism we examined changes in the proteome of LG01 and LG02 cultivated on glucose versus LG at mid-log phase. For LG01, the largest and most significant changes we observed for LG growth were an increase in the abundance of 19 proteins encoded in two distinct regions in the genome of LG01 (**Figures 2a, 3a**). For LG02, the largest and most significant changes were observed in 12 proteins encoded in three genomic regions (**Figures 2b, 3b**). The first genomic region identified in LG02 is situated at the start of a contig, suggesting it is incomplete.

**Fig. 2.**
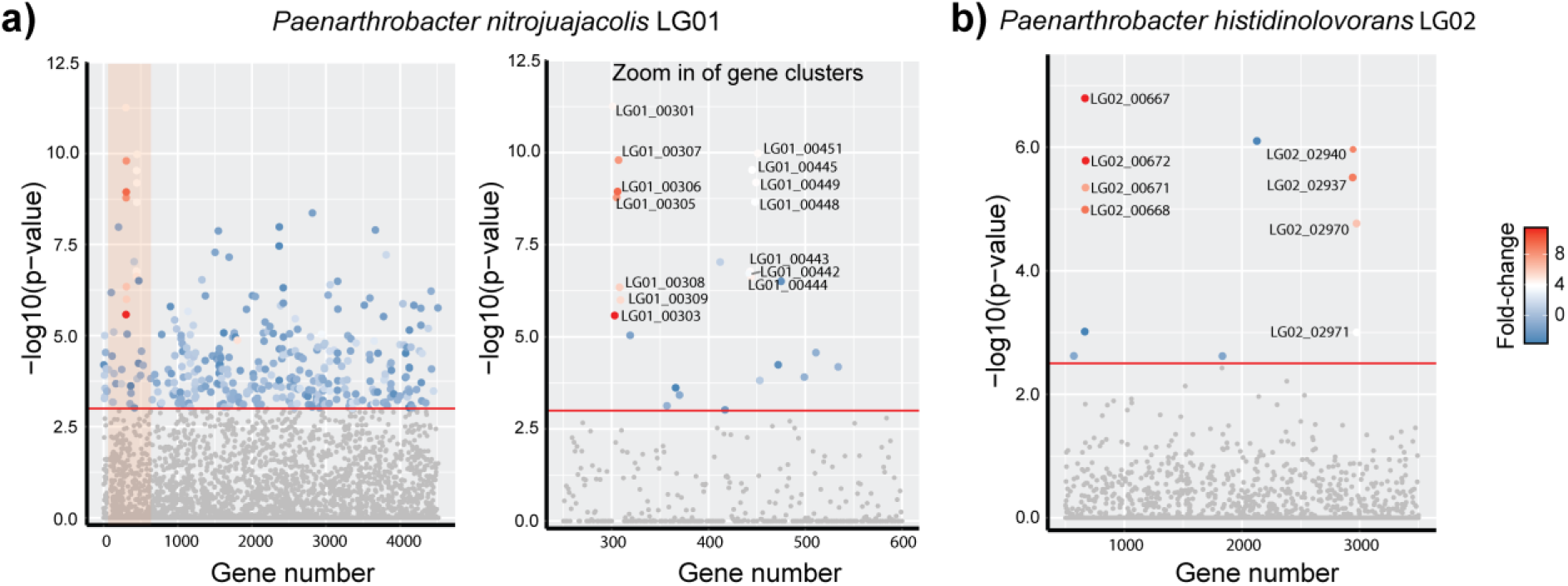
Manhattan plot of comparative proteomics data for a) *Paenarthrobacter nitrojuajacolis* LG01 and b) *Paenarthobacter histidinolovorans* LG02, grown on LG versus glucose. The relative fold change, denoted by the colour, and the −log10(p-value), denoted by on the *y*-axis, of the observed differences are plotted according to gene order, *x*-axis, revealing two regions containing up-regulated proteins in LG01 and three regions in LG02.

**Fig. 3.**
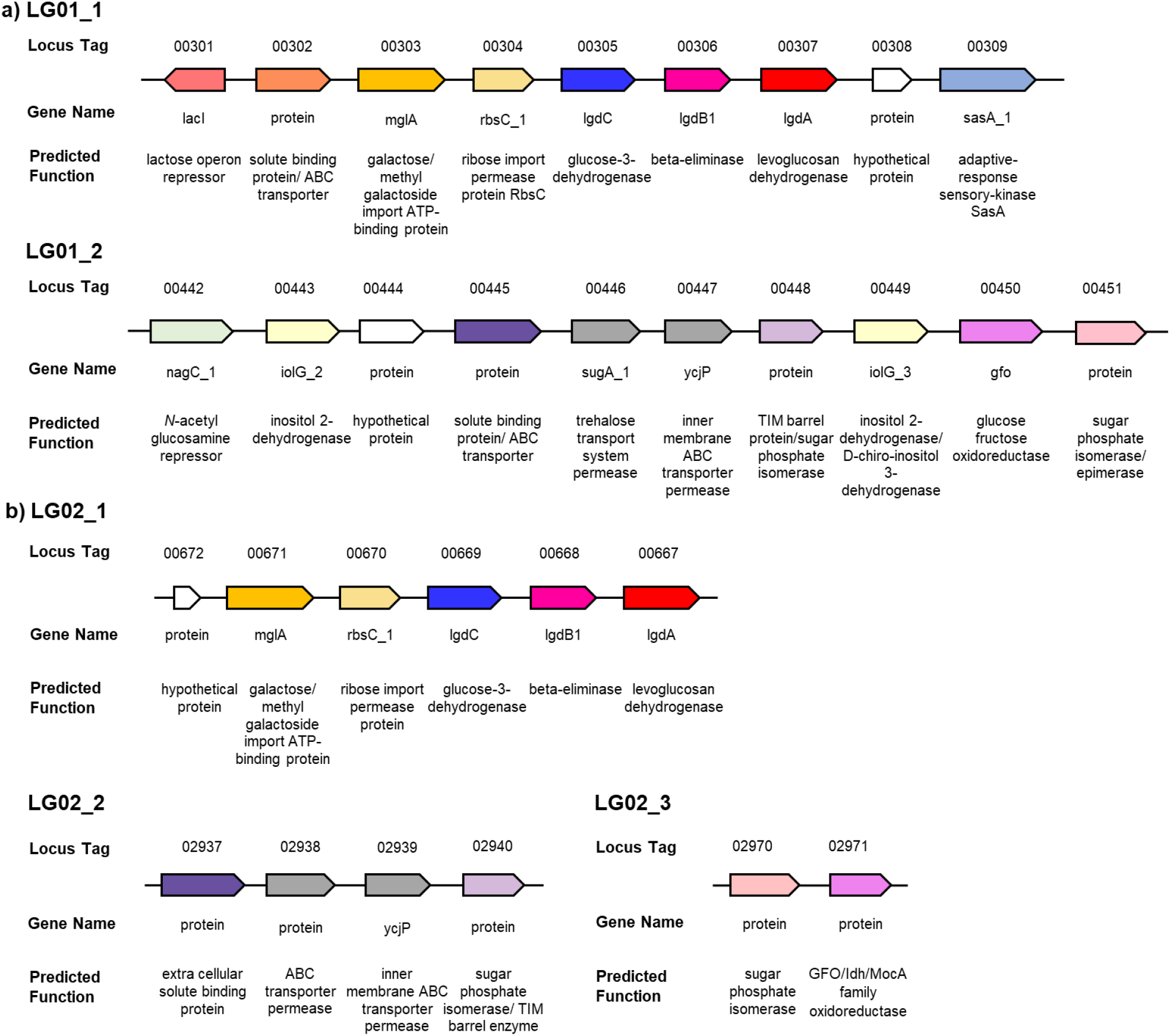
Genes identified by comparative proteomics analysis. a) Up-regulated genes identified in *Paenarthrobacter nitrojuajacolis* LG01. b) Up-regulated genes identified in *Paenarthrobacter histidinolovorans* LG02.

### SSN analysis of LG metabolizing genetic loci

The LgdA sequence from *P. nitrojuajacolis* LG01 was used as a query for a BLAST search on the phmmer webserver, which retrieved 379 sequences with *E*-value < 130. These were used to build a hidden Markov model (HMM) that was used as a query in the hmmsearch server. This retrieved 631 sequences with *E*-value < 130, which were filtered to *E*-value < 200 (percentage identity > 60%) to give 265 sequences. These sequences were used to build an SSN using the Enzyme Function Initiative tools (https://efi.igb.illinois.edu/). At an alignment score of >170, this SSN breaks into subclusters that are closely aligned with taxonomy (**Figure 4a**). The sequences were used to retrieve the genome neighbourhoods (± 10 open reading frames around *lgdA*), providing 4877 neighbours (for representative examples, see **Figure 4b**). The *lgdA* neighbours were used to build an SSN of neighbours (SSNN) (**Figure 5**, **Figure S3**).

**Fig. 4.**
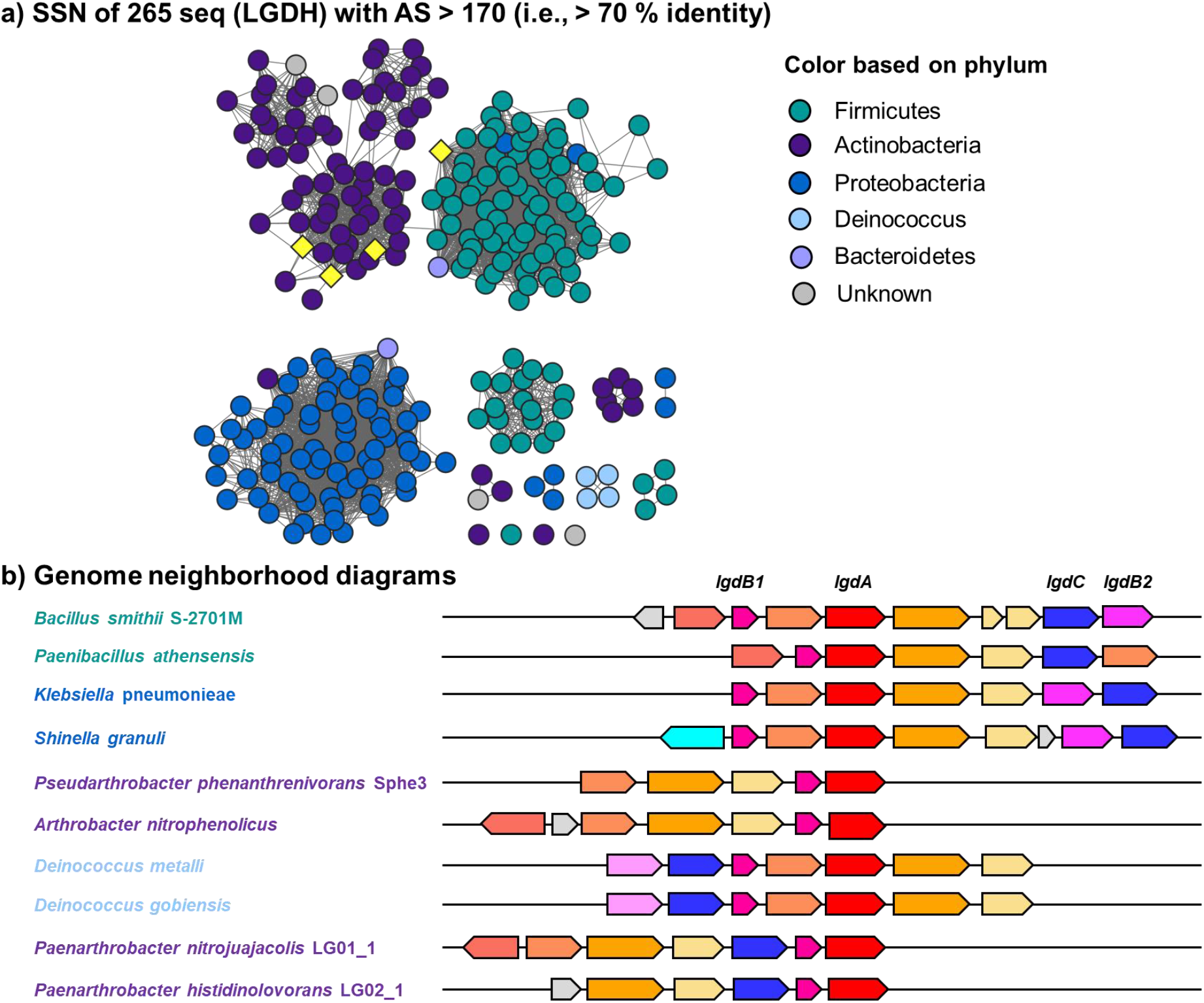
**a)** Sequence similarity network of LgdA orthologs with alignment score 170 (i.e., 70% identity). Each node is an LgdA ortholog belonging to PF01408 and PF0289, and nodes are coloured based on phylum Firmicutes, Actinobacteria, Proteobacteria, Deinococcus, Bacteroidetes and unknown. Yellow diamonds indicate proteins functionally characterized: *Pseudarthrobacter phenanthrenivorans* Sphe3 and *Bacillus smithii* S-2701M, and in this study: *Paenarthrobacter nitrojuajacolis* LG01 and *Paenarthrobacter histidinolovorans* LG02. **b)** Genome neighbourhood diagrams (GND) for different species that contains LgdA (red), LgdB1 (bright pink), LgdB2 (magenta) and LgdC (blue). Species names are coloured according to phylum as for the SSN.

**Fig. 5.**
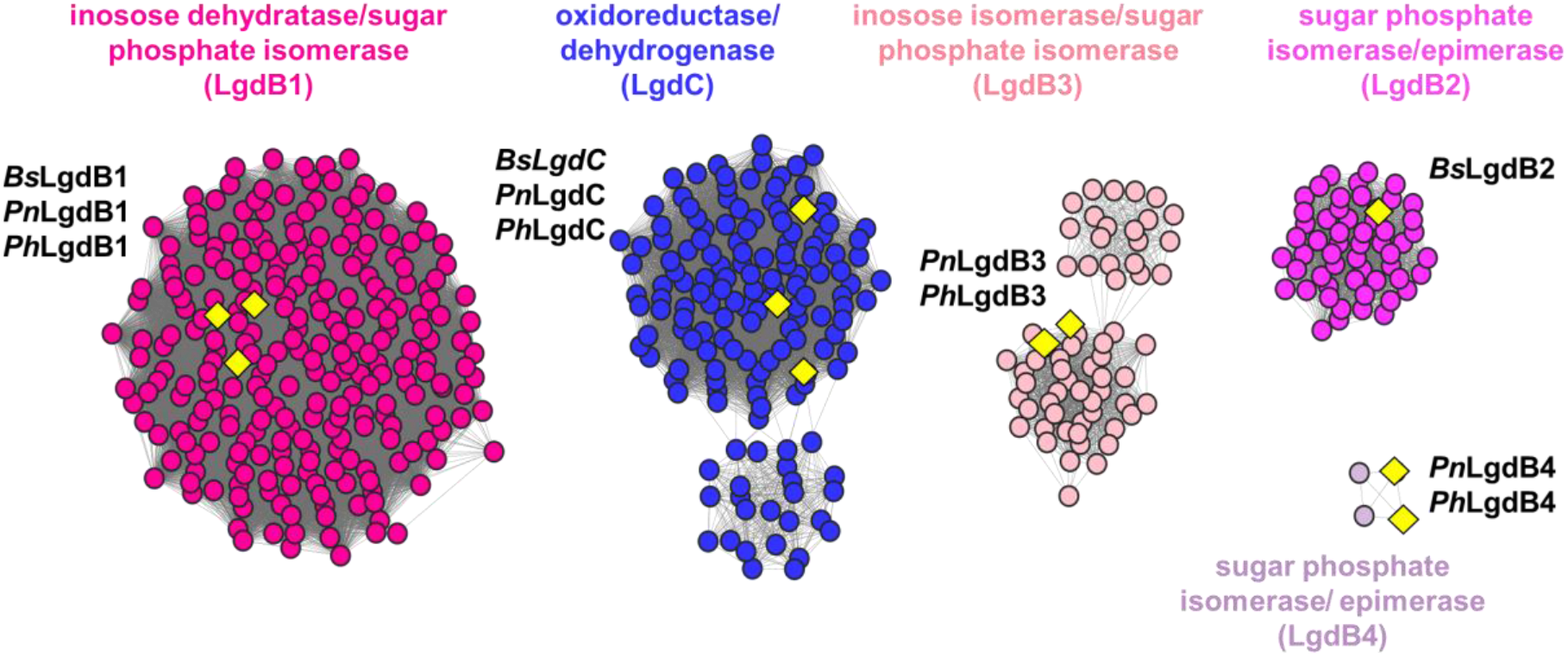
Partial sequence similarity network of neighbours (SSNN) of LgdA. The SSNN was generated with an alignment score of 50. Isofunctional clusters are coloured according to their assigned function. LgdB1 and LgdB2 map onto two clusters coloured bright pink and magenta. LgdC maps onto a blue cluster. Additional clusters annotated as inosose dehydratase/sugar phosphate isomerase (light pink) and sugar phosphate isomerase/epimerase (new one) contain proteins annotated in this work as LgdB3 and LgdB4. Yellow diamonds denote proteins that have been studied previously or in this study. From *Bacillus smithii* S-2701M: *Bs*LgdB1, *Bs*LgdB2, *Bs*LgdC; from *Paenarthrobacter nitrojuajacolis* LG01: *Pn*LgdB1, *Pn*LgdB3, *Pn*LgdC; from *Paenarthrobacter histidinolovorans* LG02: *Ph*LgdB1, *Ph*LgdB3, *Ph*LgdC. For complete SSNN see **Figure S3**.

## Discussion

Bacteria able to grow on LG were selected by sequential subculturing of media containing LG as sole carbon source and inoculated with soil. ^13^C NMR analysis revealed complete consumption of LG and no apparent carbon containing intermediates, supporting their ability to completely consume all the carbon in LG. Genome sequencing and analysis of the 16S rRNA revealed these to be *Paenarthrobacter nitrojuajacolis* LG01 and *Paenarthrobacter histidinolovorans* LG02. Comparative proteomics of samples of each bacteria grown on glucose or LG allowed identification of groups of genes that were upregulated in the LG-grown bacteria (**Figure 2**). The identified regions within both LG01 (**Figure 3a**) and LG02 (**Figure 3b**) encode close homologs of *P. phenanthenivorans* Sphe3 and *B. smithii* S-270M LGDH proteins with 96.7% and 70.1% identity (for LG01) and 97.9% and 70.7% (for LG02), respectively. Both organisms also encode homologs of LgdB1 and LgdC, but with lower percentage identity (**Tables S3 and S4**). Collectively, the presence of genes encoding LgdA, LgdB1 and LgdC provides strong evidence for canonical LGDH pathways. Neither organism contains genes encoding homologs of LgdB2. Given that the genomic region encoding LG metabolism in LG02 appears incomplete, we focused our attention on LG01.

Previous work demonstrated that *B. smithii* contains four genes *lgdA-lgdB1-lgdB2-lgdC* that between them contribute to degradation of LG (Kuritani et al. 2020), whereas Arya and co-workers identified several organisms, such as *P. athensensis* MEC069 and *Microbacterium* MEC084 that appeared to contain only *lgdA-lgdB1* (Arya et al. 2022). On the other hand, LG01 contains *lgdA-lgdB1-lgdC*. LgdA and LgdC belong to the Gfo/Idh/MocA superfamily, which contains NAD(P)-dependent oxidoreductases that act on a wide range of hydroxylated substrates (Taberman et al. 2016). LgdB1 and LgdB2 belong to protein family PF01261 within the PFAM database, a large family of proteins that contain diverse activities including inositol catabolism (Fry et al. 2001).

To gain insight into the structure and composition of LgdA-encoding genomic regions, we used SSN analysis. SSN analysis uses all-by-all pairwise sequence alignments to generate networks that show sequence relationships as a graph (Atkinson et al. 2009). We retrieved 265 sequences by an HMM search approach. The SSN was constructed by choosing an alignment score that allowed clustering based on taxonomy. Using the Enzyme Function Initiative tools (https://efi.igb.illinois.edu/), the 265 LgdA sequences were used to retrieve genome neighborhoods (± 10 ORFs) surrounding each *lgdA* gene. Examples of genome neighbourhood diagrams for bacteria from different phyla are provided in **Figure 4b**. Each of these genomic regions show conservation of *lgdB1*, but not of *lgdB2* or *lgdC* (*vide infra*).

The genomic regions surrounding each *lgdA* gene (±10 ORFs) were used to construct an SSN of neighbours (SSNN) (**Figure 5, Figure S3**). The SSNN contains isofunctional protein clusters that allow analysis of the common neighbours of *lgdA*. The SSNN reveals three large protein clusters that contain components of ABC transporters that are presumably dedicated to the importation of LG: ATP binding protein, substrate binding protein (for binding LG), and ABC transporter permease; along with several smaller clusters containing putative multifunction transporters (MFS), which presumably also facilitate binding and import of LG. Multiple clusters are present that correspond to ROK, LytTR, PucR and LacI type transcription factors.

Numerous oxidoreductase clusters are present in the SSNN. Three clusters, including the largest oxidoreductase cluster, belong to PFAM protein families PF01408 and PF02894, and correspond to the nucleotide binding and C-terminal domains of LgdC/LgdA. The LgdC sequences from *B. smithii*, and LG01 and LG02 are all located in the largest oxidoreductase cluster. Six clusters belong to PF01261, are annotated as sugar phosphate isomerase or xylose isomerase/epimerase. LgdB1 from *B. smithii*, LG01 and LG02 are present within the largest of these clusters, which contains proteins of around 275 residues. LgdB2 from *B. smithii* is located within a smaller cluster, which contains proteins of around 325 residues. The remaining four clusters do not contain any characterized members. The genomes of LG01 and LG02 encode proteins that fall into two of these orphan clusters, which we have named LgdB3 (71 sequences in cluster) and LgdB4 (4 sequences in cluster).

To visualize the organization and composition of the LG degrading gene groups we coloured the SSN of LgdA homologues according to the presence of *lgdB1, lgdB2, lgdB3* and *lgdC* in their genome neighbourhood (**Figure 6a-d**). Almost all LgdA homologs (253 out of 265) have LgdB1 encoded in their gene neighbourhood, while only a minority have LgdB2 (57 out of 265) or LgdB3 (71 out of 265) encoded in their vicinity. A Venn diagram reveals that: (1) 126 organisms contain only LgdB1; (2) LgdB2 always co-occurs with LgdB1 but never with LgdB3; and (3) LgdB3, with one exception, co-occurs with LgdB1 (**Figure 6e**). LgdB1 is widely distributed, occurring in Actinobacteria, Firmicutes, Proteobacteria, Deinococcus and Bacteroidetes. LgdB2 is more restricted, occurring mainly in Firmicutes and one Deinococcus. LgdB3 occurs Actinobacteria and Proteobacteria. Collectively, these data suggest that LgdB2 and LgdB3 may have overlapping function, but are not essential for LG metabolism. The rarer LgdB4 sugar phosphate isomerase is encoded within just 4 Actinobacteria, and co-occurs with only LgdA and LgdB1.

**Fig. 6.**
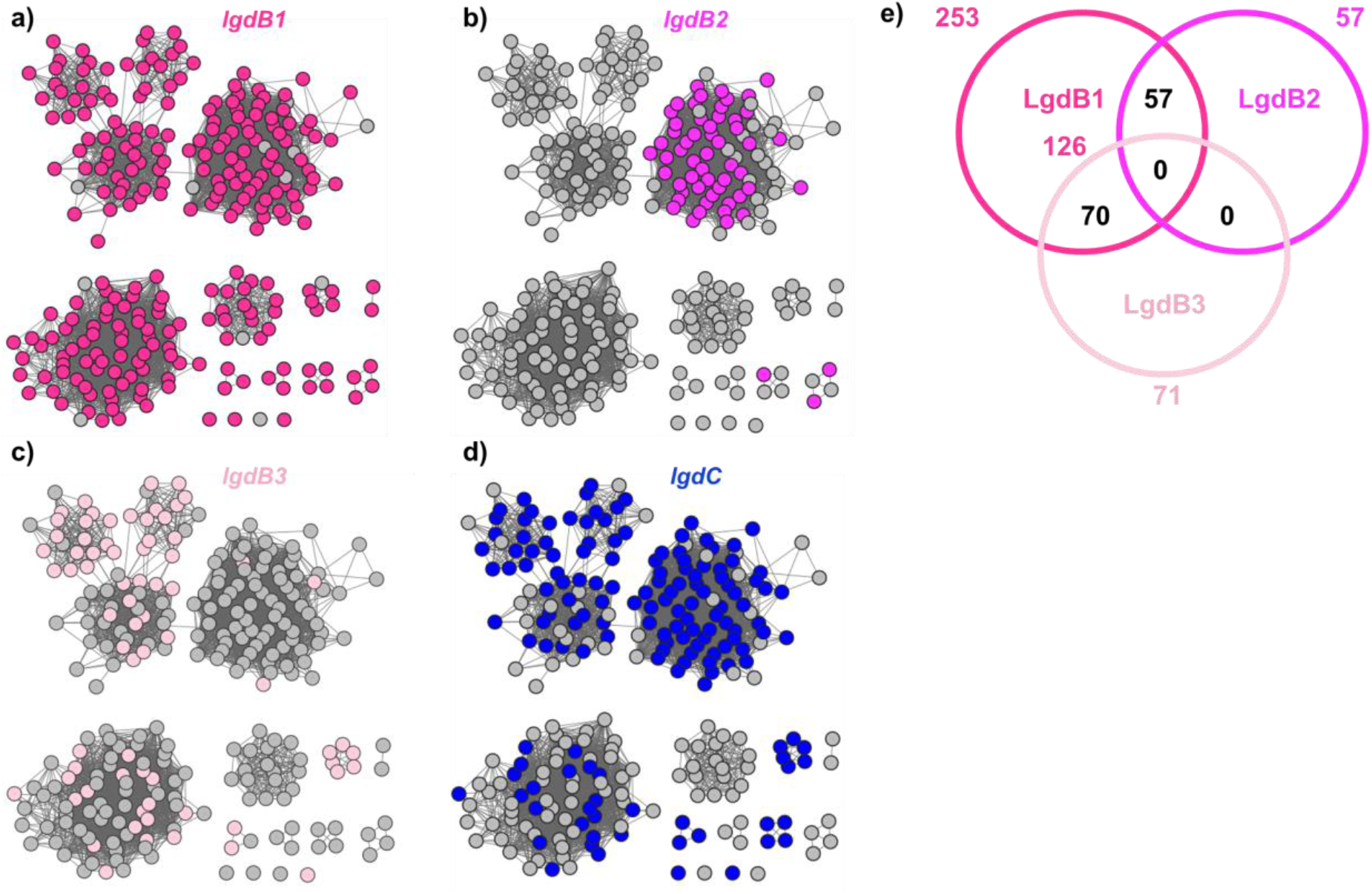
Sequence similarity network (SNN) of LgdA orthologues with alignment score 170 (i.e. 70% identity). Each node is an LgdA ortholog belong to PF01408 and PF02894. (**a-d**), nodes are coloured according to presence of specified LG degrading pathway enzymes located within an ORF window ± 10 of LgdA (each LG degrading gene belongs to a specific cluster in SSNN, **Figure 5**). LgdA nodes are coloured if they possess the following within their genome neighbourhood: a) LgdB1 (bright pink), b) LgdB2 (magenta); c) LgdB3 (light pink); d) LgdC (blue). **e)** Venn diagram shows co-occurrence of LgdB1, LgdB2 and LgdB3 in each genome neighbourhood surrounding LgdA.

Approximately half (151 out of 265) of LgdA-containing organisms have a gene for LgdC in their genome neighbourhood. It has been suggested that LgdA may possess dual activity, allowing it to function as an oxidoreductase capable of both oxidizing LG and reducing 3-keto-glucose, which could explain the absence of LgdC in some bacteria (Arya et al. 2022). LgdC is distributed across Actinobacteria, Firmicutes, Proteobacteria and Deinococcus.

Only a single 3D structure has been experimentally determined for an LG-degrading protein in the LGDH pathway, LGDH from *P. phenanthrenivorans* Sphe3 (PDB 6A3I) (Sugiura et al. 2018). Comparison of this structure with the AlphaFold2 model for LgdA from LG01 shows almost identical 3D fold (RMSD 0.252 Å; **Figure 7a**), and the presence of nucleotide cofactor and LG binding sites (**Figure 7b**). A comparison of the AlphaFold2 models for *B. smithii* and LG01 LgdB1 proteins shows similar 3D folds (RMSD 0.433 Å) (**Figure 7c**), suggestive of similar function. The peptide chain of LgdB1 is much shorter than LgdB2, LgdB3 and LgdB4, and the comparisons of AlphaFold models showed low similarity and high RMSD values (**Table S4b**). Overlay of *B. smithii* LgdB2, LG01 LgdB3 (RMSD 1.345 Å versus *B. smithii* LgdB2), and LgdB4 (RMSD 3.288 Å versus *B. smithii* LgdB2) shows broadly similar (αβ)_8_ barrel folds, but weaker structural similarity (**Figure 7d**). The weaker structural similarity is difficult to interpret in the absence of structural studies to determine the molecular basis of substrate recognition and catalysis, but suggests that these proteins could have a similar function, may act on modified intermediates in the pathway (eg 6-phospho-2-hydroxy-3-keto-d-glucal, formed by hexokinase), or may not be involved in the pathway.

**Figure 7.**
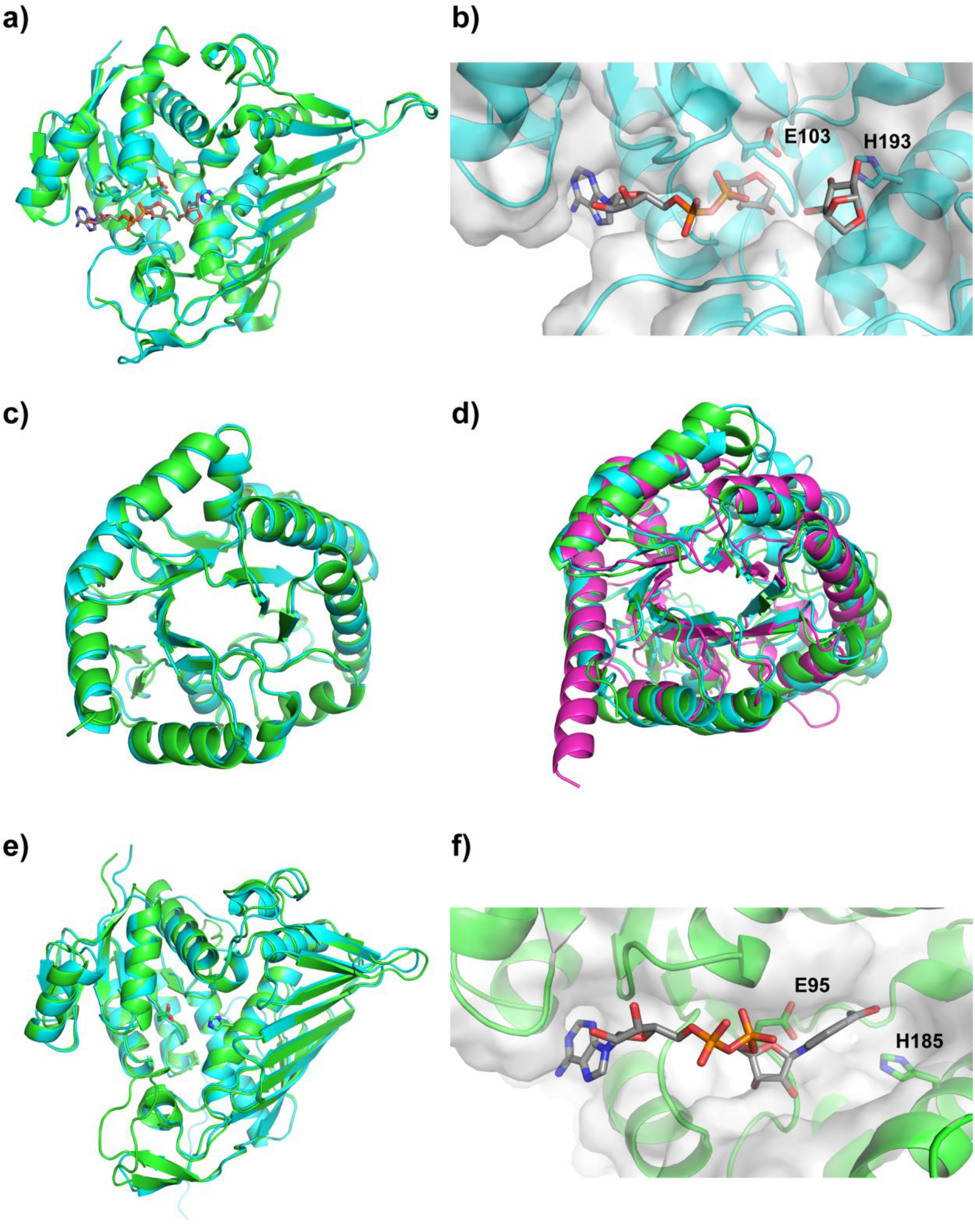
LG01 Alphafold2 models and overlays with experimentally determined structure of LgdA. (a) An overlay of the model for LgdA from LG01 (cyan) with levoglucosan dehydrogenase from *Pseudarthrobacter phenanthrenivorans* Sphe3 (green, PDB ID: 6A3I). (b) Close-up view of the putative active site of LgdA from LG01 (cyan) with nucleotide cofactor and levoglucosan substrate overlaid from 6A3I, and the proposed catalytic residues E103 and H193 shown in stick form. (c) An overlay of the models for LG01 and *B. smithii* LgdB1 (cyan and green, respectively). (d) An overlay of the models for *B. smithii* LgdB2 (green), LG01 LgdB3 (cyan) and LG01 LgdB4 (magenta). (e) An overlay of the models for *B. smithii* and LG01 LgdC (cyan and green, respectively). (f) A close-up view of the putative active site of LG01 LgdC, with nucleotide cofactor overlaid from 6A3G, and the putative catalytic residues E95 and H185 shown in stick form.a

A comparison of the AlphaFold2 models of *B. smithii* and LG01 LgdC reveals similar 3D folds (RMSD 0.641 Å) (**Figure 7e**). Overlay of LG01 LgdC and the *P. phenanthrenivorans* LgdA in complex with NADH gave a model that showed LgdC could accommodate NADH, consistent with its demonstrated function as an NADH-dependent glucose 3-dehydrogenase (**Figure 7f**).

## Conclusions

Two LG degrading soil bacteria belonging to the *Paenarthrobacter* genus (strains LG01 and LG02) were isolated by enrichment culture involving growth on LG. These organisms effect the complete degradation of LG, and give a stationary phase optical density on LG that is similar to that on glucose. Comparative proteomics revealed the presence of key genes encoding homologues of proteins involved in LG degradation, namely LGDH (LgdA), 3-keto-LG β-eliminase (LgdB1), and glucose 3-dehydrogenase (LgdC). However, the upregulated proteins included several additional members of the PFAM PF01261 sugar phosphate isomerase/xylose isomerase family that exhibit low homology to LgdB1. Structural modelling shows that these adopt similar folds to LgdB1 and their occurrence within the genomic regions of a wide range of bacteria encoding *lgdA-lgdB1-lgdC* genes suggest that they may assist in the processing of intermediates in the LG-degradation pathway.

## Supporting information

Supplementary material file 2, Accession codes

## Abbreviations

ABC: ATP-binding cassette
DNA: Deoxyribonucleic acid
GND: Genome neighbourhood diagram
HMM: Hidden Markov Model
LG: Levoglucosan
LGDH: Levoglucosan dehydrogenase
NADH: reduced nicotinamide adenine dinucleotide
ORF: open reading frame
SSN: Sequence similarity network
SSNN: Sequence similarity network of neighbours

## Authors’ contributions

SJW conceived the study with substantial input from AK. AK performed enrichment culture and microbial growth and characterization. NS conducted proteomics. MH and SP performed genome sequencing, assembly and analysis. AK and EGB performed bioinformatic analysis. AK and SJW wrote the paper with input and approval of the final draft from the authors.

## Availability of data and material

The genome datasets generated and/or analysed during the current study are available in the NCBI database with BioProject accession codes PRJNA925556 (strain LG01) and PRJNA925557 (strain LG02). The mass spectrometry proteomics data has been deposited in the Proteome Xchange Consortium via the PRIDE partner repository with the data set identifier: PXD039086.

## Code availability

Not applicable.

## Declarations

### Ethics approval

Not applicable

### Consent to participate

Not applicable

### Consent for publication

Not applicable

