## Supplementary material file 1 for "Identification of levoglucosan degradation pathways in bacteria and sequence similarity network analysis"

import pandas as pd

import sqlite3

con=sqlite3.connect("Data/LGDH_neighbors_output.sqlite")

df = pd.read_sql_query("SELECT accession from neighbors", con)

con.close()

df.to_csv('Data/LGDH_neighbors.list', header=False, index=False)

**Figure S1.** Python script used to extract the accession codes of the retrieved neighbors from the SQlite file downloaded from EFI-GNT tools.

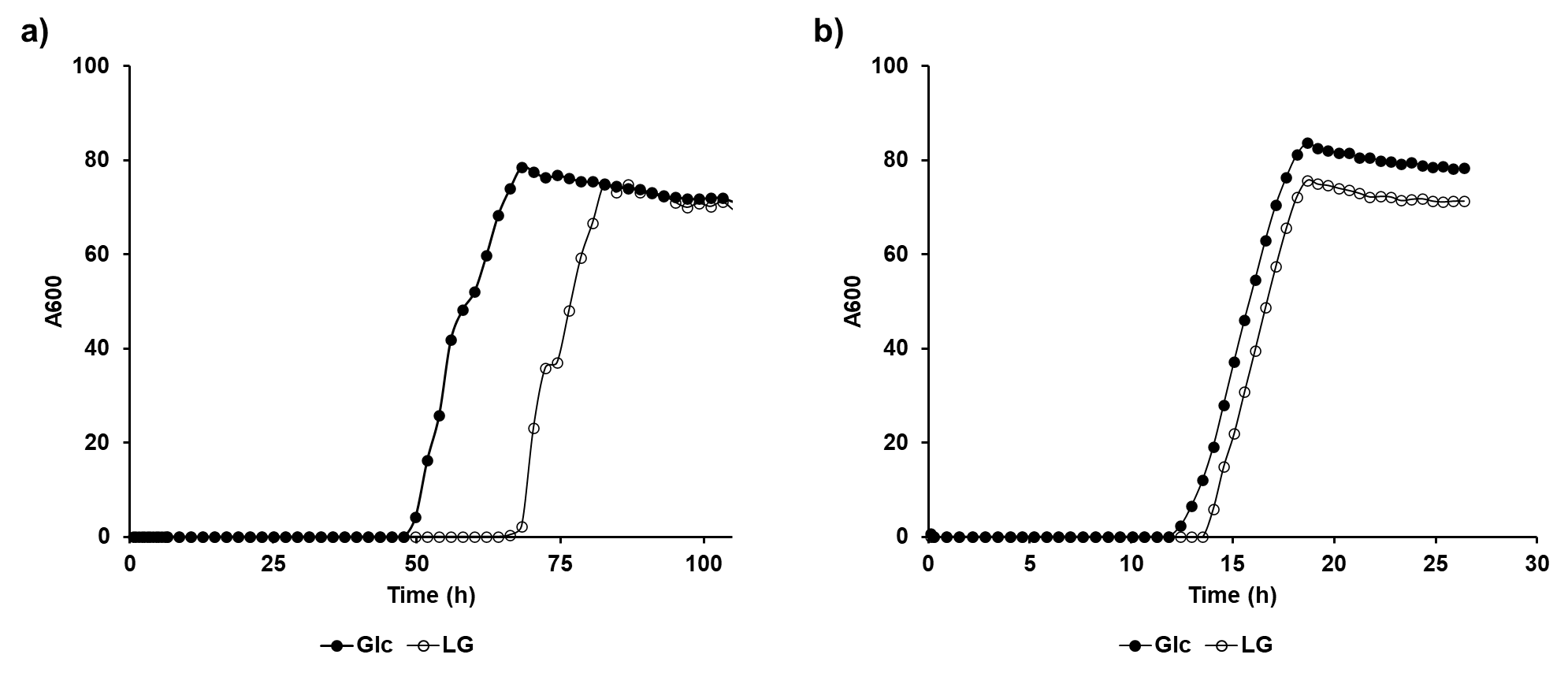

**Figure S2.** Growth curves of *Paenarthrobacter* strains a) LG01 and b) LG02 grown on M9 media containing 5 mM glucose or SQ.

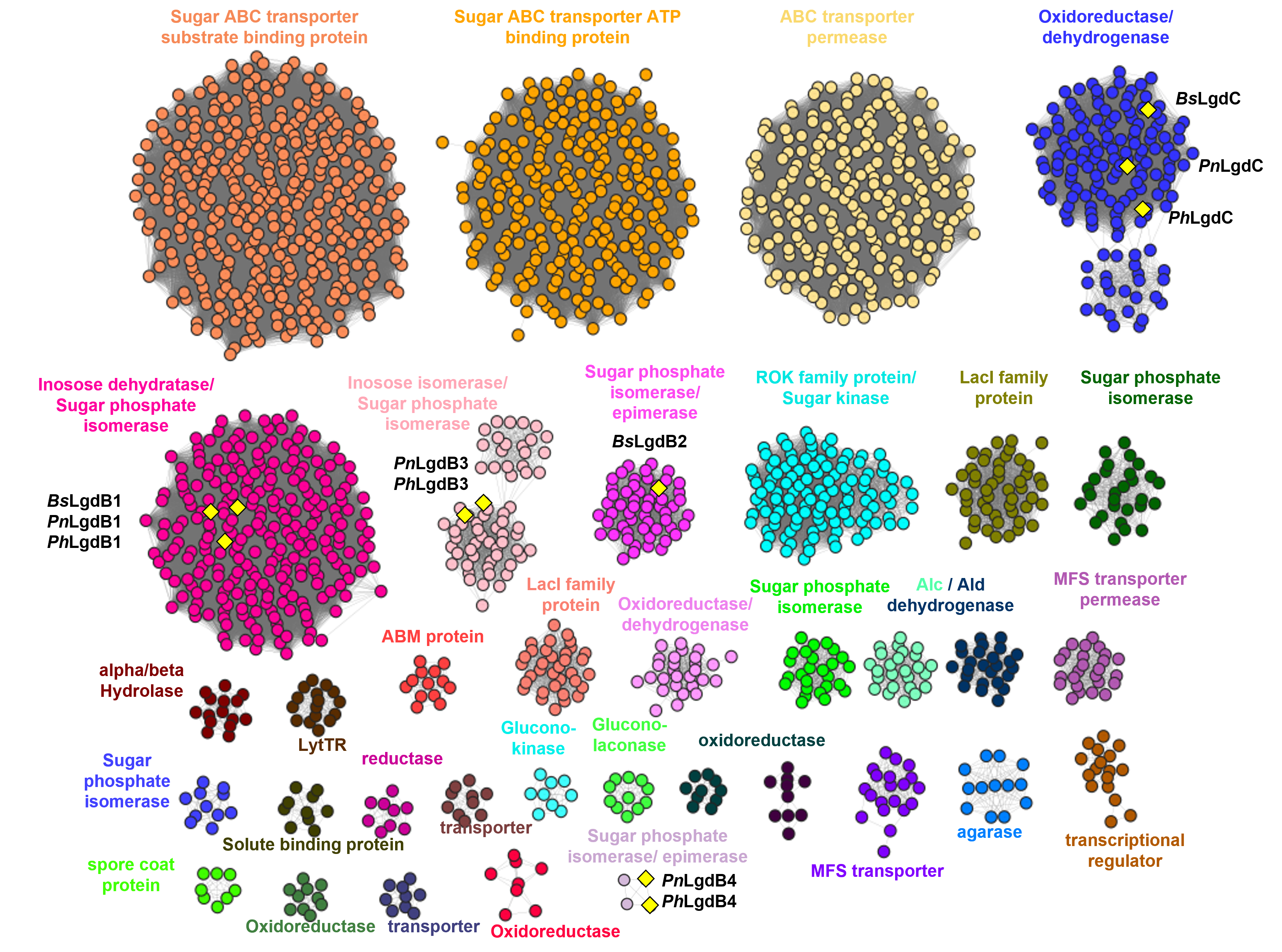

**Figure S3.** Sequence similarity network of neighbours (SSNN) of LgdA. SSNN was generated with an alignment score of 50. Isofunctional clusters are coloured accordingly to their assigned function. Three large clusters are for ABC transporters (orange, yellow and light yellow). LgdB1 and LgdB2 map onto two clusters coloured bright pink and magenta. LgdC maps onto a blue cluster. Additional clusters annotated as inosose dehydratase/sugar phosphate isomerase (light pink) and sugar phosphate isomerase/epimerase (new one) contain proteins annotated in this work as LgdB3 and LgdB4. Yellow diamonds denote proteins that have been studied previously or in this study. From *Bacillus smithii* S-2701M: *Bs*LgdB1, *Bs*LgdB2, *Bs*LgdC; from *Paenarthrobacter nitrojuajacolis* LG01: *Pn*LgdB1, *Pn*LgdB3, *Pn*LgdC; from *Paenarthrobacter histidinolovorans* LG02: *Ph*LgdB1, *Ph*LgdB3, *Ph*LgdC. Clusters smaller than 7 members are not shown, except for the cluster for LgdB4.

**Table S1. Classification and general feature of *Paenarthrobacter* spp. LG01 and LG02**

| **MIGS ID** | **Property** | **Term** | **Evidence code** |
| --- | --- | --- | --- |
|  | Classification | Domain *Bacteria* | TAS [1] |
|  |  | Phylum *Actinobacteria* | TAS [2] |
|  |  | Class *Actinomycetia* | TAS [3] |
|  |  | Order *Micrococcales* | TAS [3-6] |
|  |  | Family *Micrococcaceae* | TAS [3-5, 7] |
|  |  | Genus *Paenarthrobacter* | TAS [8] |
|  |  | Species *Paenarthrobacter nitrojuajacolis* LG01 | TAS [8] |
|  |  | Species *Paenarthrobacter histidinolovorans* LG02 | IDA |
|  | Gram Strain | Not measured |  |
|  | Cell Shape | Not reported |  |
|  | Motility | Not reported |  |
|  | Sporulation | Not reported |  |
|  | Optimum Temperature | Not tested, used 30℃ | IDA |
|  | pH range; Optimum | Not tested, used 7-8 | IDA |
|  | Carbon source | Yeast extract/tryptone, glucose, levoglucosan | IDA |
| MIGS-6 | Habitat | Soil | IDA |
| MIGS-22 | Oxygen requirement | Aerobic | IDA |
| MIGS-15 | Biotic relationship | Free living | IDA |
| MIGS-4 | Geographic location | Melbourne, VIC, Australia | IDA |
| MIGS-5 | Sample collection | June 11, 2021 | IDA |
| MIGS-4.1 | Latitude | -37.7949428 | IDA |
| MIGS-4.2 | Longitude | 144.9525146 | IDA |
| MIGS-4.4 | Altitude | Not reported |  |

*Evidence codes - IDA: Inferred from Direct Assay; TAS: Traceable Author Statement (i.e., a direct report exists in the literature)

**Table S2. Table of accession codes for levoglucosan degrading genes.**

| **Annotation** | **Description** | **Accession code** |
| --- | --- | --- |
| ***Bs*LgdA** | levoglucosan dehydrogenase | BCB28822.1 |
| ***Bs*LgdB1** | β-eliminase | BCB28820.1 |
| ***Bs*LgdB2** | 3-keto-glucose-dehydratase | BCB28827.1 |
| ***Bs*LgdC** | glucose-3-dehydrogenase | BCB28826.1 |
| ***Pp*LgdA** | levoglucosan dehydrogenase | ADX72256.1 |

*Bs* = *Bacillus smithii* S-2701M, *Pp* = *Pseudarthobacter phenanthrenivorans* Sphe3.

**Table S3. Percentage identity for LG degrading proteins: Oxidoreductases LgdA and LgdC.**

|  | ***Pp*LgdA** | ***Bs*LgdA** | ***Pn*LgdA** | ***Ph*LgdA** | ***Pn*LgdC** | ***Ph*LgdC** |
| --- | --- | --- | --- | --- | --- | --- |
| ***Pp*LgdA** |  | **70.1** | **96.7** | **97.9** | 33.3 | 33.3 |
| ***Bs*LgdA** |  |  | **71.5** | **70.7** | 32.2 | 31.9 |
| ***Pn*LgdA** |  |  |  | **97.4** | 32.3 | 32.9 |
| ***Ph*LgdA** |  |  |  |  | 32.9 | 32.9 |
| ***Pn*LgdC** |  |  |  |  |  | **96.9** |
| ***Ph*LgdC** |  |  |  |  |  |  |

Bold shows percentage identity >50%. *Pp* = *Pseudarthrobacter phenanthrenivorans* Sphe3, *Bs* = *Bacillus smithii* S-2701M, *Pn* = *Paenarthrobacter nitrojuajacolis* LG01 and *Ph* = *Paenarthrobacter histidinolovorans*

**Table S4. Percentage identity for LG degrading proteins: LgdB1, LgdB2, LgdB3, LgdB4 (annotated as sugar phosphate isomerase).**

|  | ***Bs*LgdB1** | ***Bs*LgdB2** | ***Pn*LgdB1** | ***Pn*LgdB3** | ***Pn*LgdB4** | ***Ph*LgdB1** | ***Ph*LgdB3** | ***Ph*LgdB4** |
| --- | --- | --- | --- | --- | --- | --- | --- | --- |
| ***Bs*LgdB1** |  | 31.7 | **55.5** | 22.0 | 27.6 | **55.6** | 25.4 | 30.5 |
| ***Bs*LgdB2** |  |  | 24 | 26.7 | - | 23.0 | 26.4 | **-** |
| ***Pn*LgdB1** |  |  |  | - | 25.6 | **97.4** | 23.4 | 25.2 |
| ***Pn*LgdB3** |  |  |  |  | 23.2 | 26.1 | **97.6** | 24.0 |
| ***Pn*LgdB4** |  |  |  |  |  | 29.9 | 22.8 | **91.5** |
| ***Ph*LgdB1** |  |  |  |  |  |  | 22.7 | 29.9 |
| ***Ph*LgdB3** |  |  |  |  |  |  |  | - |
| ***Ph*LgdB4** |  |  |  |  |  |  |  |  |

Bold shows percentage identity >50%. *Pp* = *Pseudarthrobacter phenanthrenivorans* Sphe3, *Bs* = *Bacillus smithii* S-2701M, *Pn* = *Paenarthrobacter nitrojuajacolis* LG01 and *Ph* = *Paenarthrobacter histidinolovorans.*

**Table S5. RMSD values for structural comparison: Oxidoreductases LgdA and LgdC.**

|  | ***Pp*LgdA** | ***Pn*LgdA** | ***Bs*LgdC** | ***Pn*LgdC** |
| --- | --- | --- | --- | --- |
| ***Pp*LgdA** |  | 0.252 | 0.869 | 0.963 |
| ***Pn*LgdA** |  |  | 0.882 | 0.994 |
| ***Bs*LgdC** |  |  |  | 0.641 |
| ***Pn*LgdC** |  |  |  |  |

**Table S6. RMSD values for structural comparison: LgdB1, LgdB2, LgdB3, LgdB4 (annotated as sugar phosphate isomerase).**

|  | ***Bs*LgdB1** | ***Bs*LgdB2** | ***Pn*LgdB1** | ***Pn*LgdB3** | ***Pn*LgdB4** |
| --- | --- | --- | --- | --- | --- |
| ***Bs*LgdB1** |  | 2.705 | 0.433 | 5.326 | 2.218 |
| ***Bs*LgdB2** |  |  | 2.782 | 1.345 | 3.288 |
| ***Pn*LgdB1** |  |  |  | 11.361 | 1.443 |
| ***Pn*LgdB3** |  |  |  |  | 7.613 |
| ***Pn*LgdB4** |  |  |  |  |  |
